# *Tg*mRHel drives unified RNA processing of *coxI* mRNA generated from a complex mitochondrial genomic context

**DOI:** 10.64898/2026.05.29.728635

**Authors:** Zala Gluhic, Sabrina Tetzlaff, Peter Liebers, Nikiforos Drakoulis, Alexandra Possling, Capella S. Maguire, Victor Flores, Ross F. Waller, Giel G. van Dooren, Christian Schmitz-Linneweber

## Abstract

The mitochondrial genome of *Toxoplasma gondii* is highly fragmented and recombination-prone, creating a structurally dynamic genetic landscape. How such a genome is used efficiently to produce functional mRNAs remains unclear: it is unknown whether transcription draws from many alternative genomic configurations or a restricted subset, and how any resulting precursor RNAs are processed into mature transcripts. More broadly, mitochondrial RNA processing mechanisms in this system are poorly understood.

Here, we show that recombination of the *T. gondii* mitochondrial genome generates a diverse population of structurally distinct precursor RNAs for the essential cytochrome c oxidase subunit I (*coxI*) protein. Rather than being derived from a single defined primary transcript, these heterogeneous precursors are unified into a single mature mRNA through a post-transcriptional mechanism dependent on the DEAD-box RNA helicase *Tg*mRHel. *Tg*mRHel is required for 5′-end processing of *coxI* mRNA and for the accumulation of mitochondrial rRNAs, both of which are essential for Complex IV biogenesis, oxidative phosphorylation, and parasite survival. Loss of *Tg*mRHel leads to the accumulation of diverse *coxI* precursor RNAs and a failure to generate the mature transcript.

Our findings reveal an unexpected genome–transcriptome interface in which extensive genomic variability is not suppressed at the DNA level but instead resolved through RNA helicase–mediated processing. This work establishes a new conceptual framework for how gene expression fidelity can be maintained in the context of highly dynamic, recombining genomes.

## Introduction

Apicomplexan mitochondria are indispensable for parasite viability. They support essential cellular processes such as ATP production, iron–sulfur cluster biosynthesis and pyrimidine biosynthesis [1]. Disruption of mitochondrial function is therefore lethal in both *Plasmodium* and *Toxoplasma*, as demonstrated by pharmacological inhibition of the mitochondrial electron transport chain [ETC; 2,3]. Most ETC proteins are imported from the cytosol, but cytochrome c oxidase subunits I and III (CoxI and CoxIII), and cytochrome b (Cob) are encoded in the mitochondrial genome. Consistent with this, both pharmacological and genetic inhibition of mitochondrial gene expression have been shown to reduce ETC function and arrest parasite development [4–11].

Mitochondrial gene expression in apicomplexan parasites represents a striking example of evolutionary innovation and reduction. Unlike the relatively conserved mitochondrial genomes of most eukaryotes, those of apicomplexans - such as *Plasmodium* and *Toxoplasma* - are highly atypical: linear, fragmented, and extremely reduced in gene content with most encoding only the three ETC components mentioned above [12–15]. Paralleling this genomic minimalism is a set of unconventional RNA biology features, including extensive rRNA fragmentation, an unusual mitoribosome structure and non-canonical translation mechanisms [9,10,13,16]. These features suggest that RNA maturation in apicomplexan mitochondria follows a highly divergent pathway. Starting from long precursor RNAs [14,17], RNA processing down to rRNA fragments and mature mRNAs is known to take place, but how this is achieved and which factors are involved is currently unclear.

In many species, RNA helicases play central roles in the regulation of gene expression within mitochondria [18]. RNAs frequently form stable secondary structures, including stem-loops and other folded domains, particularly in intron-containing pre-mRNAs and untranslated regions. These structures can pose physical barriers to essential RNA processing events such as splicing, cleavage, and editing, and can also hinder the initiation of translation by obstructing ribosome binding. RNA helicases use the energy from ATP hydrolysis to unwind or remodel these structures, thereby enabling access by the appropriate enzymatic or ribonucleoprotein complexes. A well-studied example is the DEAD-box RNA helicase Mss116p in yeast mitochondria, which facilitates the splicing of group I and II introns through RNA structure remodeling [19,20].

Beyond facilitating RNA processing, helicases are also required for the assembly and maintenance of functional mitochondrial ribosomes. Several helicases contribute to the proper folding of rRNAs and the assembly of ribosomal subunits, which are essential for efficient translation of mitochondrial-encoded proteins [21,22]. For instance, the DEAD-box RNA helicase DDX28 in human mitochondria is necessary for the maturation of the large ribosomal subunit; its absence results in the accumulation of incomplete or dysfunctional ribosomes, with direct consequences for organellar protein synthesis (Tu and Barrientos 2015) .

Additionally, RNA helicases are integral components of mitochondrial RNA surveillance and decay pathways. The maintenance of mitochondrial transcriptome integrity depends on the selective degradation of aberrant, misprocessed, or excess RNAs. Helicases such as SUV3, in complex with exonucleases like PNPase, unwind structured RNA substrates to enable their efficient degradation [23]. This function is critical for preventing the accumulation of non-functional or potentially deleterious RNAs, and for regulating gene expression post-transcriptionally. Collectively, RNA helicases act at multiple levels of RNA metabolism to ensure that mitochondrial gene expression is efficient, accurate, and responsive to cellular needs.

In *Toxoplasma gondii*, the mitochondrial genome is highly fragmented and organized into a series of short, non-redundant sequence blocks that are maintained and transcribed as linear concatemers [13,14]. These concatemers give rise to unusually long precursor RNAs that contain multiple gene segments [14]. The dynamic, recombining nature of the *T. gondii* mitochondrial genome is expected to lead to a high degree of transcriptomic complexity, with the potential generation of numerous distinct precursor RNAs differing in gene content and arrangement. This poses a significant challenge for RNA processing, as coding sequences for mRNAs and rRNAs must be accurately identified, excised, and matured from heterogeneous transcript pools. To date, the molecular machinery responsible for processing these precursor RNAs into functional mRNAs and rRNAs remains entirely unknown. Here we demonstrate that a mitochondrial RNA helicase, *Tg*mRHel (*Toxoplasma gondii* mitochondrial RNA helicase), has an essential role in the maturation of RNA transcripts in *T*. *gondii* mitochondria, providing functional insights into gene expression from the highly fragmented mitochondrial genome of *T*. *gondii* parasites.

## Results

### *Tg*mRHel is a mitochondria-localized DEAD-box helicase essential for parasite growth

Given the importance of RNA helicases in mitochondrial RNA metabolism, we systematically analyzed all genes annotated as RNA helicase in the *T. gondii* genome for their subcellular localization using DeepLoc [S1 Table, 24]. Amongst the 85 proteins annotated as helicases in the *T. gondii* genome, three were predicted to be targeted to mitochondria and had previously been detected in mitochondrial proteomic datasets [LOPIT, 25]. Of these candidates, the gene TGGT1_313240 exhibited the strongest negative fitness score in a genome-wide CRISPR screen [26], suggesting an important role for parasite survival. This gene is annotated as a DEAD-box helicase and was therefore designated *T. gondii* mitochondrial RNA Helicase (*Tg*mRHel).

Sequence analysis revealed that *Tg*mRHel contains the canonical DEAD motif and all typical sequence features characteristic of the DDX5/Dbp2 subfamily of DEAD-box helicases (Fig 1A, S1A). Members of this subfamily include the well-characterized helicases DDX17 and DDX5 in metazoans, although these proteins are not mitochondrial [27]. Structural prediction indicates that *Tg*mRHel adopts the typical DEAD-box helicase architecture (S1A, B Fig). Unlike its metazoan homologs, *Tg*mRHel possesses an N-terminal extension of ∼340 amino acids predicted to be largely intrinsically disordered (S1B, C Fig). Phylogenetic analysis showed that *Tg*mRHel clusters with related DEAD-box helicases from other diverse alveolate lineages (S1D Fig), all of which are also predicted to localize to mitochondria. The closest homolog within *T. gondii* is TGME49_236650, an orthologue of DDX5 (S1D Fig) that is not predicted to localize to mitochondria. *Tg*mRHel most likely originated from an ancient gene duplication event from an ancestral DDX5-like helicase (S1D Fig). Together, these analyses support the hypothesis that *Tg*mRHel is a mitochondrial DEAD-box helicase.

**Fig 1.**
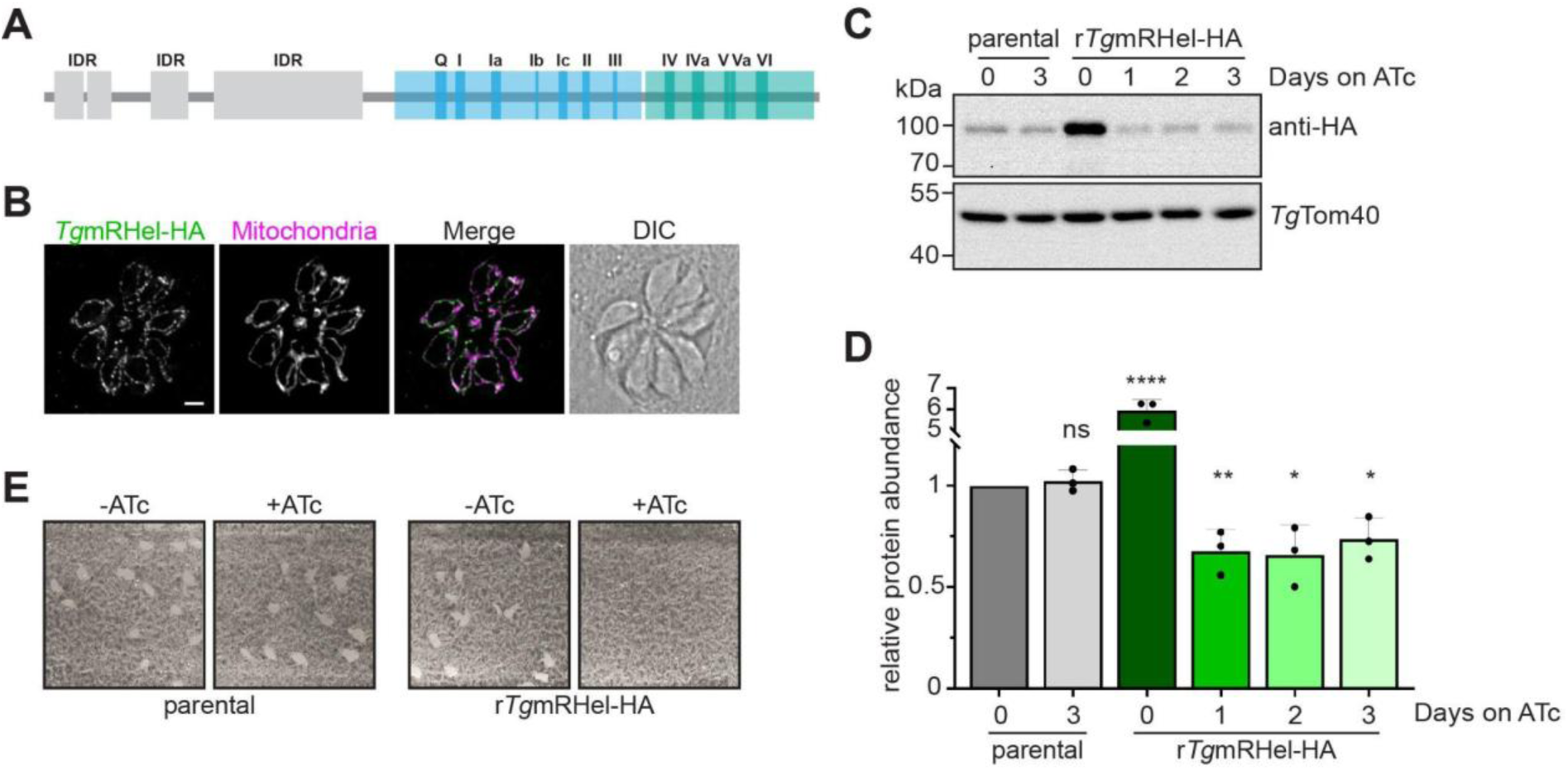
Induced depletion of *Tg*mRHel leads to reduced parasite growth. A. Domain structure of the *Tg*mRHel DEAD-box RNA helicase. All canonical DEAD RNA helicase domains are found in the primary sequence of *Tg*mRHel. Domains 1 and 2 are marked in blue and green, respectively. In addition to the conserved domains, *Tg*mRHel contains a number of sequence motifs that are recognized as intrinsically disordered domains (IDRs). B. Immunofluorescence assay of an eight-cell vacuole of *Tg*mRHel-HA parasites probed with anti-HA (green) and anti-*Tg*Tom40 antibodies to label the mitochondrion (magenta). Scale bar is 2 µm. C. Western blots of proteins extracted from r*Tg*mRHel-HA parasites cultured in the absence of ATc or in the presence of ATc for 1–3 days next to the parental line *Tg*mRHel-HA cultured without or with ATc for three days. Samples were separated by SDS-PAGE, and probed with anti-HA and anti-*Tg*Tom40 (loading control) antibodies. D. Quantification of Western blot results (C) normalised to *Tg*mRHel-HA protein abundance in parental parasite strain cultured without ATc. Unpaired t-tests were performed to compare each condition to this untreated parental reference and asterisks indicate statistical significance according to predefined p-value thresholds (*p < 0.05, **p < 0.01, ****p < 0.0001, ns = not significant, n = 3). E. Plaque assays measuring proliferation of *Tg*mRHel-HA and r*Tg*mRHel-HA parasites cultured in the absence (left) or presence (right) of ATc for 8 days. Assays are from a single experiment and are representative of three independent experiments.

To experimentally test the localization of *Tg*mRHel, we introduced a C-terminal haemagglutinin (HA) epitope tag into the endogenous locus, generating the *Tg*mRHel-HA parasite line (S2A, B Fig). Immunofluorescence assays using anti-HA antibodies revealed a signal that co-localized with the mitochondrial marker *Tg*Tom40, demonstrating that *Tg*mRHel localizes to the mitochondrion (Fig 1B).

To investigate the function of *Tg*mRHel, we generated a conditional knockdown parasite line by replacing the endogenous promoter of *Tg*mRHel-HA with an anhydrotetracycline (ATc)-regulated promoter (S2C, D Fig). This line, termed regulatable *Tg*mRHel-HA (r*Tg*mRHel-HA) allows inducible repression upon ATc addition. Western blot analysis showed that *Tg*mRHel protein levels were constitutively upregulated in this line (Fig 1C). After induction, they were strongly reduced within one day of ATc treatment and remained low during continued exposure (Fig 1C). In contrast, *Tg*mRHel protein levels remained unchanged in the parental *Tg*mRHel-HA line treated with ATc for three days (Fig 1C). Although *Tg*mRHel was not completely eliminated in the knockdown line, quantification confirmed a substantial reduction in protein abundance after ATc treatment (Fig 1D). Furthermore, these results indicate that the ATc-inducible promoter is significantly stronger than the native *Tg*mRHel promoter. Overall, these results demonstrate that *Tg*mRHel expression can be efficiently downregulated using this system.

To determine whether *Tg*mRHel is required for parasite proliferation, we compared growth of parental *Tg*mRHel-HA and r*Tg*mRHel-HA parasites in plaque assays. Parasites were cultured in the absence or presence of ATc for eight days. While parental parasites formed plaques irrespective of ATc treatment, r*Tg*mRHel-HA parasites failed to form plaques in the presence of ATc, indicating a severe defect in parasite proliferation upon *Tg*mRHel depletion (Fig 1E). These findings demonstrate that normal *Tg*mRHel levels are essential for the growth of tachyzoites.

To confirm that the growth defect is specifically caused by *Tg*mRHel depletion, we generated complementation lines expressing an additional copy of *Tg*mRHel driven by an ATc-insensitive alpha-tubulin promoter and fused to a C-terminal Ty1 epitope (c*Tg*mRHel-Ty1/r*Tg*mRHel-HA). Three independent clones (G10, E7, D6) were analyzed. Western blotting showed efficient downregulation of the HA-tagged endogenous allele upon ATc treatment, while the Ty1-tagged copy remained stably expressed (S3A Fig). Consistent with this, all complementation clones retained normal plaque formation in the presence of ATc, whereas the r*Tg*mRHel-HA knockdown line failed to grow under these conditions (S3B Fig). These results demonstrate that the proliferation defect is specifically due to loss of *Tg*mRHel and that expression of a copy fully rescues parasite growth.

Together, these results establish that *Tg*mRHel is a mitochondrially localized DEAD-box RNA helicase that is essential for the proliferation of *T. gondii*.

### Reduced oxygen consumption and loss of Complex III and IV following downregulation of *Tg*mRHel

Because mitochondrial gene expression is essential for the production of key subunits of the mitochondrial ETC, we next tested whether depletion of *Tg*mRHel affects mitochondrial respiration. To assess ETC activity, we measured the basal mitochondrial oxygen consumption rate (mOCR) in extracellular parasites using a Seahorse extracellular flux assay [28]. Parental *Tg*mRHel-HA and r*Tg*mRHel-HA parasites were cultured either in the absence or presence of ATc for 1–3 days prior to analysis. Depletion of *Tg*mRHel resulted in a rapid decline in basal mOCR, reaching a reduction of more than 95% after two days of ATc treatment (Fig 2A). In contrast, ATc treatment had no effect on mOCR in parental parasites (Fig 2A), indicating that the reduction in respiration is specifically caused by *Tg*mRHel depletion.

**Fig 2.**
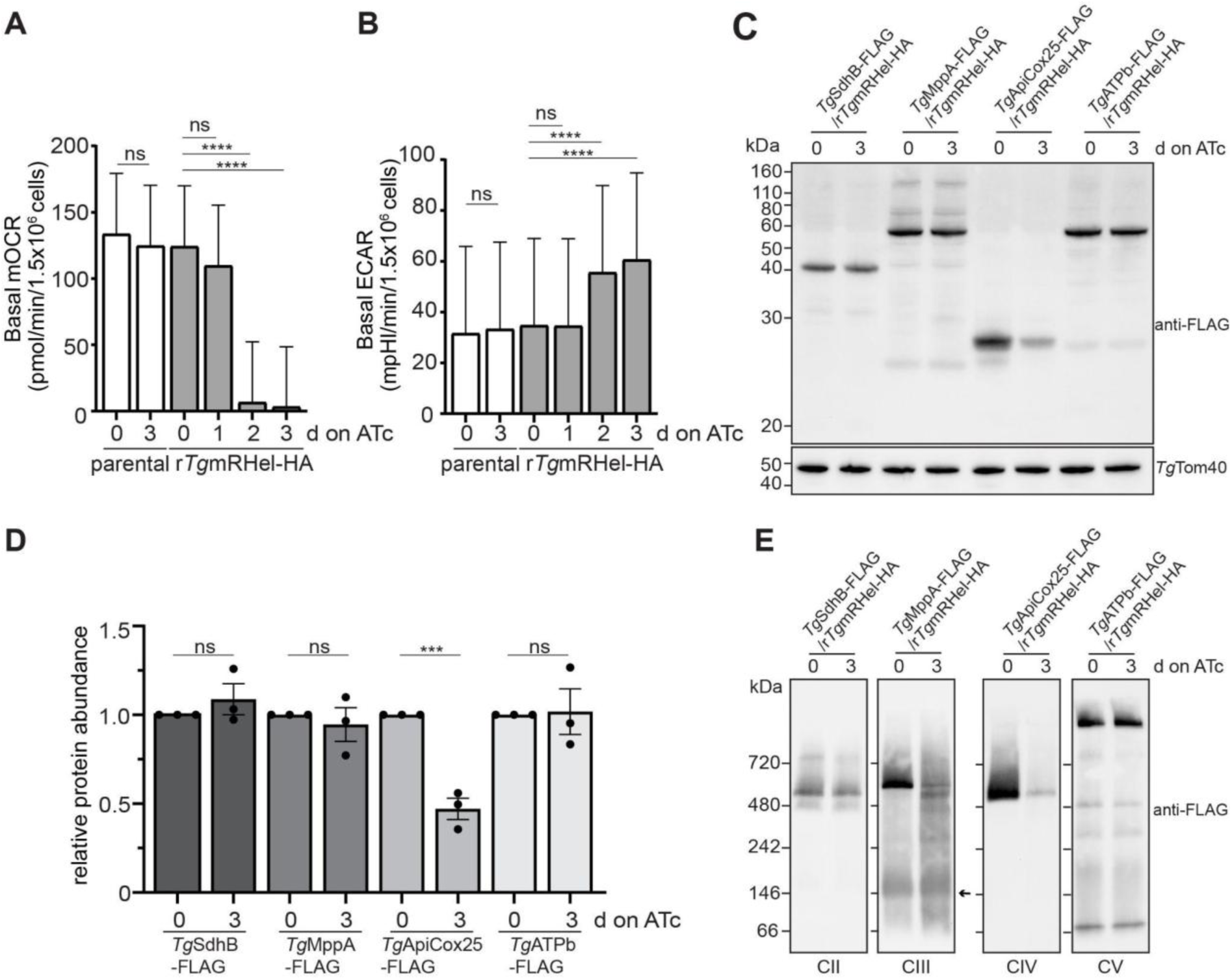
Characterization of OxPhos capacity after *Tg*mRHel depletion. A. Basal mitochondrial O_2_ consumption rate (mOCR) of parental *Tg*mRHel-HA (white) and r*Tg*mRHel-HA (grey) parasites cultured in the absence of ATc or in the presence of ATc for 1–3 days. A linear mixed-effects model was fitted to the data, and values depict the estimated marginal mean ± 95% CI of three independent experiments. One-way ANOVA followed by Tukey’s multiple pairwise comparisons test was performed and asterisks indicate statistical significance according to predefined p-value thresholds (****p < 0.0001, ns = not significant). B. Basal extracellular acidification rate (ECAR) of parental (white) and r*Tg*mRHel-HA (grey) parasites cultured in the absence of ATc or in the presence of ATc for 1–3 days. A linear mixed-effects model was fitted to the data, and values depict the estimated marginal mean ± 95% CI of three independent experiments. One-way ANOVA followed by Tukey’s multiple pairwise comparisons test was performed and asterisks indicate statistical significance according to predefined p-value thresholds (****p < 0.0001, ns = not significant). C. Immunoblot analysis of proteins extracted from *Tg*SdhB-FLAG/r*Tg*mRHel-HA, *Tg*MppA-FLAG/r*Tg*mRHel-HA, *Tg*ApiCox25-FLAG/r*Tg*mRHel-HA and *Tg*ATPb-FLAG/r*Tg*mRHel-HA parasites cultured without ATc or with ATc for three days. Samples were separated by SDS-PAGE, and probed with anti-FLAG and anti-*Tg*Tom40 antibodies. Immunoblots are representative of three biological replicates. D. Quantification of Western blot results (C) normalised to protein abundance in parasite strain cultured without ATc. Unpaired t-test was performed and asterisks indicate statistical significance according to predefined p-value thresholds (***p < 0.001, ns = not significant, n = 3). E. Imunoblot analysis of proteins extracted from *Tg*SdhB-FLAG/r*Tg*mRHel-HA, *Tg*MppA-FLAG/r*Tg*mRHel-HA, *Tg*ApiCox25-FLAG/r*Tg*mRHel-HA and *Tg*ATPb-FLAG/r*Tg*mRHel-HA parasites cultured without or for three days with ATc. Samples were prepared in 1% digitonin for *Tg*SdhB-FLAG and *Tg*ATPb-FLAG and 1% (v/v) TX-100 for *Tg*MppA-FLAG and *Tg*ApiCox25-FLAG, separated by BN-PAGE, and detected with anti-FLAG antibody. Arrow indicates the putative MPP complex.

A severe reduction in oxygen consumption could reflect either a specific defect in mitochondrial respiration or a general loss of cell viability. To distinguish between these possibilities, we measured the basal Extracellular Acidification Rate (ECAR), which reflects proton release associated with glycolysis. r*Tg*mRHel-HA parasites maintained measurable basal ECAR following ATc treatment and in fact showed increased basal ECAR relative to untreated controls (Fig 2B). Such an increase is consistent with a metabolic shift toward glycolysis when oxidative phosphorylation is impaired. These results indicate that parasites remain metabolically active after *Tg*mRHel depletion and that the observed respiratory defect is not a consequence of cell death.

Since the three protein-coding genes in the *T. gondii* mitochondrial genome encode subunits of Complex III and IV, and since these may depend on *Tg*mRHel activity for their expression, we next assessed the impact of *Tg*mRHel knockdown on the accumulation of the complexes of the ETC. We generated r*Tg*mRHel-HA parasite lines expressing FLAG-tagged versions of the Complex III subunit *Tg*MppA and the Complex IV subunit *Tg*ApiCox25. As controls, we also generated parasite lines expressing FLAG-tagged versions of the Complex II subunit *Tg*SdhB and the ATP synthase (Complex V) subunit *Tg*ATPb, both of which belong to complexes composed exclusively of nuclear-encoded proteins (S4A,B,D,F,H Fig). All four tagged proteins localized to the mitochondrion as expected (S4C,E,G,I Fig). Parasites expressing these tagged proteins were cultured in the absence or presence of ATc and analyzed by SDS-PAGE and immunoblotting. Following *Tg*mRHel depletion, levels of *Tg*ApiCox25 were strongly reduced, whereas the abundance of *Tg*MppA, *Tg*SdhB, and *Tg*ATPb remained largely unchanged (Fig 2C,D).

Although protein levels of the nuclear Complex III subunit *Tg*MppA were unaffected, we could not exclude that Complex III might be affected as well given that the mitochondrial genome encodes its cytochrome b (Cob) subunit. Therefore, we assessed the integrity of ETC complexes by Blue Native-PAGE (BN-PAGE). This analysis confirmed that depletion of *Tg*mRHel resulted in a pronounced loss of the ∼600 kDa *Tg*ApiCox25-FLAG–containing Complex IV [Fig 2E; ,29]. In addition, the ∼675 kDa *Tg*MppA-containing Complex III was strongly diminished, indicating destabilization of Complex III. *Tg*MppA is known to participate in two distinct protein complexes: as a core subunits of Complex III and as a component of the mitochondrial processing peptidase (MPP), which cleaves mitochondrial targeting sequences from imported proteins [1,30]. We indeed additionally detected *Tg*MppA in a ∼220 kDa lower-molecular-weight complex which remained unaffected upon *Tg*mRHel depletion [Fig 2E; ,29]). The unchanged total *Tg*MppA levels observed by SDS-PAGE therefore reflect its presence in this stable complex, despite the loss of functional Complex III [30].

In contrast, the ∼500 kDa Complex II (*Tg*SdhB-FLAG) and the ∼900 kDa ATP synthase Complex V (*Tg*ATPb-FLAG) were largely unaffected by *Tg*mRHel depletion (Fig 2E). Because these complexes do not contain mitochondrially encoded subunits, their stability is independent of mitochondrial gene expression.

Together, these results demonstrate that *Tg*mRHel is required for mitochondrial respiratory activity in *T. gondii.* Loss of *Tg*mRHel leads to a rapid collapse of oxygen consumption and specifically compromises the integrity of ETC complexes that contain mitochondrially encoded subunits.

### *Tg*mRHel is required for mitochondrial rRNA accumulation

DEAD-box RNA helicases frequently participate in ribosome biogenesis [21]. To determine whether *Tg*mRHel contributed to mitochondrial ribosome formation in *T. gondii*, we examined the abundance and sedimentation behaviour of mitochondrial rRNAs following depletion of the helicase.

We first analyzed mitochondrial ribosome complexes by sucrose density gradient centrifugation (polysome analysis). Crude organelle-enriched extracts from r*Tg*mRHel-HA parasites treated without ATc or with ATc for 1–3 days were fractionated on sucrose gradients, and RNA from the fractions was analyzed by RNA gel blot hybridization. Probes specific for the mitochondrial rRNA fragments RNA3 (large ribosomal subunit) and SSUA (small ribosomal subunit) were used, both of which previously produced robust signals in such analyses (Tetzlaff et al. 2024).

Two major effects were observed. First, the overall abundance of both RNA3 and SSUA decreased markedly following *Tg*mRHel depletion (Fig 3A,B). Signal intensities progressively declined over the three-day ATc treatment period, indicating that mitochondrial rRNAs fail to accumulate when *Tg*mRHel levels are reduced. Second, the sedimentation profiles of these rRNAs changed substantially. In untreated parasites, RNA3 and SSUA signals were predominantly detected in heavier gradient fractions (fractions 10–15), corresponding to assembled ribosomal complexes. Upon *Tg*mRHel depletion, however, the remaining rRNA signals shifted toward lighter fractions (fractions 6–9), consistent with a loss of assembled ribosomes and possibly the accumulation of incomplete ribosomal assemblies (Fig 3A,B). This fractionation pattern is consistent with previous analyses of *T. gondii* mitochondrial ribosomes, in which small and large ribosomal subunits sediment in intermediate fractions and assembled monosomes appear in heavier fractions [9,13]. Quantification of the signals confirmed the redistribution of rRNAs toward lighter fractions in the knockdown parasites (Fig 3A,B).

**Fig 3.**
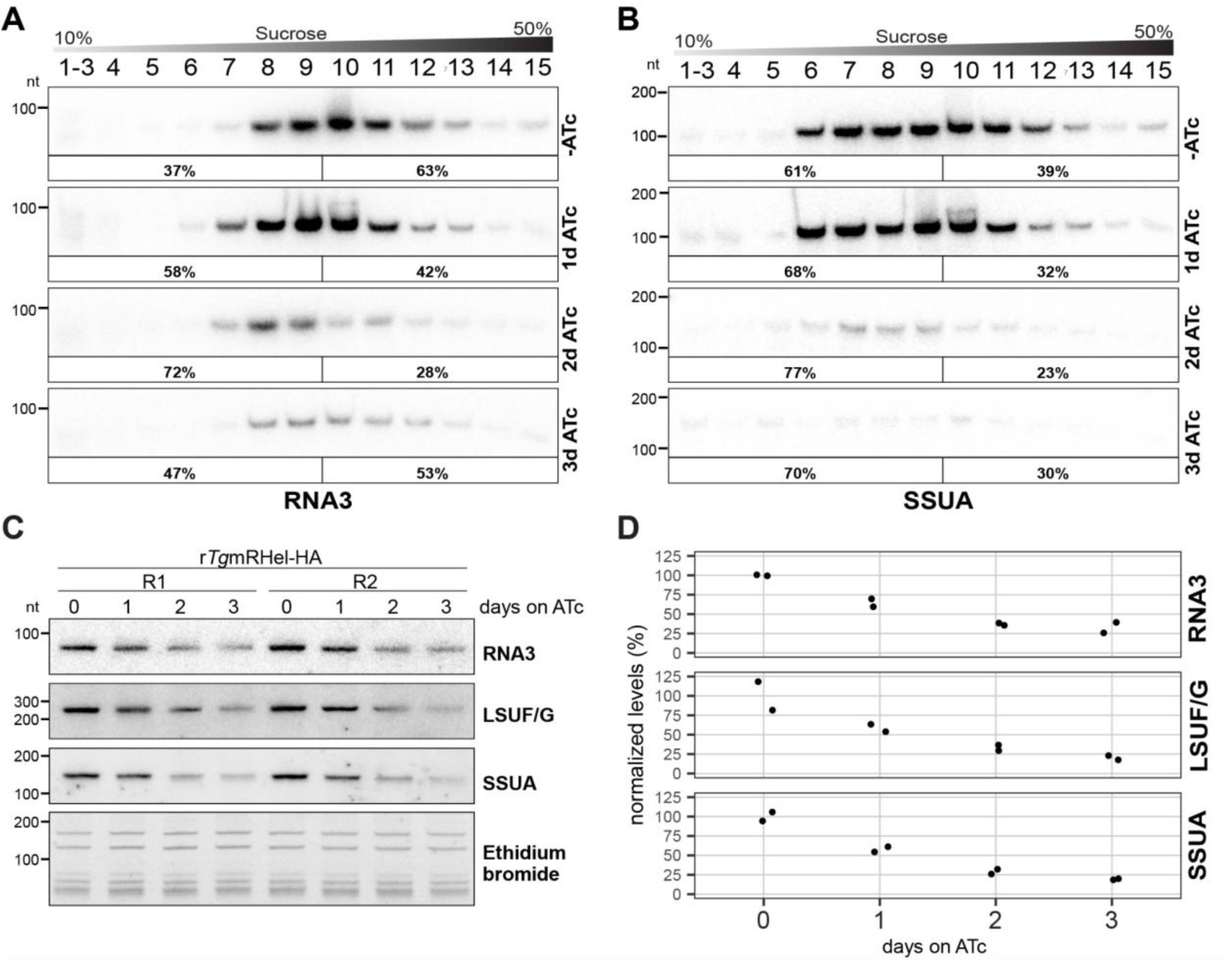
Mitochondrial rRNAs show reduced accumulation and reduced association with mono- and polysomes. A. Organelle-enriched extracts from *T. gondii* r*Tg*mRHel parasites treated with ATc for 1 to 3 days, or left untreated, were fractionated by sucrose density gradient centrifugation and analyzed by RNA gel blot hybridization. A schematic above the gel images illustrates the sucrose gradient. The panels below show hybridization results from RNA blots derived from the ethidium bromide-stained gels presented in S5 Fig using a probe against RNA3. Percentages in square brackets below each blot indicate the proportion of signal present in fractions 1–9 relative to fractions 10–15. B. After probing for RNA3, the blots were stripped using a denaturing buffer and re-hybridized with a probe targeting SSUA. C. RNA gel blot analysis of rRNA before and after *Tg*mRHel depletion. Total cellular RNA (350 ng) was run on a denaturing polyacrylamide gel and detected using the indicated ^32^P-end-labeled oligonucleotide probes after transfer to a nylon membrane. Hybridizations were performed sequentially, with probes stripped between each hybridization. Hybridization signals of the expected size are marked by arrowheads. As a loading control, an ethidium bromide stain of the gel prior to blotting is shown below the autoradiographs. D. Quantification of signals in (C) normalized to the cytosolic 5.8S rRNA signal in the ethidium bromide stain. Individual data points represent two independent repeats. For each transcript, values are visualized relative to the mean of the untreated control (0d ATc treatment), which was set to 100%.

To determine whether these effects were specific to mitochondrial ribosomes, we examined the distribution of cytosolic rRNAs in the same gradient fractions by ethidium bromide staining. Cytosolic rRNA profiles remained largely unchanged following *Tg*mRHel depletion, indicating that the observed changes are specific to mitochondrial ribosomes (S5 Fig).

The strong reduction of mitochondrial rRNA fragments in the polysome analysis prompted us to examine rRNA accumulation directly in total cellular RNA. Total RNA from r*Tg*mRHel-HA parasites cultured in the absence of ATc or the presence of ATc for 1–3 days was analyzed by RNA gel blot hybridization using probes specific for RNA3, LSUF/G (large subunit fragment) and SSUA rRNAs. All mitochondrial rRNA fragments showed reduced accumulation after one day of ATc treatment, followed by progressive loss over the subsequent days (Fig 3C,D). In contrast, cytosolic small rRNAs and tRNAs visualized by ethidium bromide staining remained unchanged, confirming equal RNA loading (Fig 3C).

Together, these results demonstrate that *Tg*mRHel is required for the accumulation and/or stability of mitochondrial rRNA for the formation of polysomes.

### Depletion of *Tg*mRHel leads to accumulation of *coxI* precursor transcripts

Reduced polysome formation can result from defects in ribosome assembly, but can also arise from impaired mRNA availability or reduced mRNA translatability. To determine whether *Tg*mRHel depletion affects mitochondrial mRNA levels, we examined the accumulation and processing of the three mitochondrially encoded mRNAs - *coxI*, *coxIII*, and *cob* - by RNA gel blot hybridization. Total RNA was isolated from r*Tg*mRHel-HA parasites treated with ATc over a three-day time course and compared to untreated controls.

The three mitochondrial mRNAs showed distinct responses to *Tg*mRHel depletion. The *cob* mRNA exhibited only a minor reduction in abundance over the time course (Fig 4A). In contrast, the *coxIII* mRNA was strongly reduced already after one day of ATc treatment (Fig 4B).

**Fig 4.**
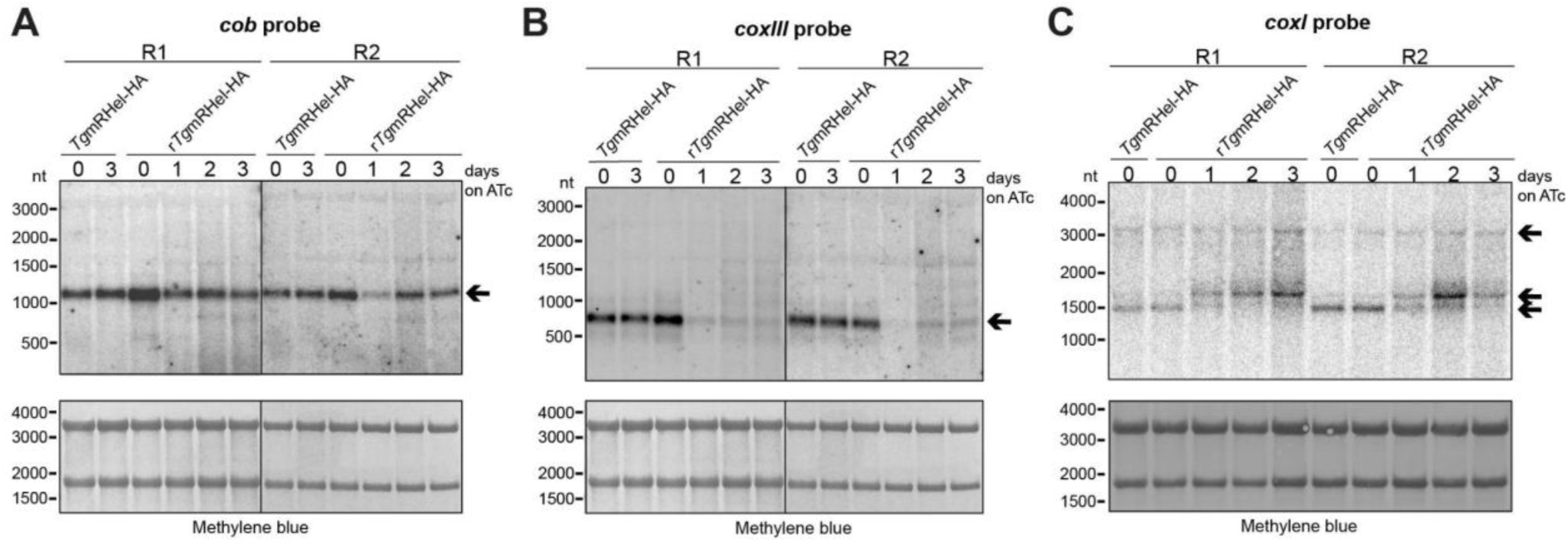
Changes in mitochondrial mRNA levels during depletion of *Tg*mRHel. A. RNA gel blot hybridizations of mitochondrial *cob* mRNA after depletion of *Tg*mRHel. Top panel: Total cellular RNA (3.85 μg per lane) from r*Tg*mRHel parasites treated with ATc for 1 to 3 days, or left untreated, was separated by agarose gel electrophoresis and transferred to a nylon membrane. The membrane was hybridized with a radiolabeled strand-specific *cob* RNA probe. Bottom panel: Cytosolic rRNAs on the same membrane were visualized by methylene blue staining and serve as a loading control. The arrow indicates the main *cob* transcript. R1 and R2 indicate independent replicates 1 and 2, respectively. B. The blot shown in (B) was stripped using a denaturing buffer and reprobed with a strand-specific radiolabeled probe against *coxIII* mRNA. The arrow indicates the main *coxIII* transcript. The same methylene blue labelled loading control as (A) is included to facilitate comparison. C. RNA gel blot analysis of *coxI* mRNA, run on a separate gel and transferred independently. The membrane was hybridized with a strand-specific radiolabeled probe against *coxI* mRNA. Cytosolic rRNAs stained with methylene blue serve as a loading control. The lower arrow indicates the main 1.5 kb *coxI* transcript. The middle arrow indicates the 1.7 kb precursor found to increase in the mutant. The unchanged 3.3 kb band may either be a longer precursor or a cross-hybridization of the probe.

The most intriguing changes were observed for *coxI* transcripts. In untreated r*Tg*mRHel-HA parasites and in the parental line, two RNA species were detected: a ∼3.3 kb transcript and a ∼1.5 kb transcript (Fig 4C). The size of the latter closely matches the expected length of the *coxI* coding sequence (1476 nucleotides), so is likely the mature *coxI* mRNA. Upon depletion of *Tg*mRHel, an additional *coxI*-specific RNA species of approximately 1.7 kb became visible as early as one day after ATc addition. Although this transcript could be faintly detected in untreated parasites, it accumulated strongly during *Tg*mRHel depletion and became the dominant *coxI* RNA species by days two and three (Fig 4C). At the same time, the mature ∼1.5 kb *coxI* mRNA progressively decreased in abundance and was nearly undetectable after three days of ATc treatment. In contrast, the signal at 3.3 kb remained largely unchanged throughout the time course. Whether this signal is indeed a long precursor RNA of *coxI* cannot be decided at present.

Taken together, these results indicate that depletion of *Tg*mRHel leads to the accumulation of an intermediate *coxI* transcript and a concomitant loss of the mature mRNA. These observations suggest that *Tg*mRHel is required for a processing step that converts precursor *coxI* transcripts into the mature mRNA. To further characterize these precursor transcripts, we next mapped the 5′ ends of *coxI* RNAs using 5′ RACE.

### *Tg*mRHel is required for 5′-end processing of *coxI* mRNA from alternative precursor transcripts

The accumulation of an intermediate *coxI* transcript upon *Tg*mRHel depletion suggested a defect in *coxI* mRNA processing. To determine the origin of these precursor transcripts and to identify the 5′ ends of *coxI* RNAs, we performed 5′ rapid amplification of cDNA ends (5′ RACE). A RACE adaptor was ligated to the 5′ ends of total RNA isolated from parental *Tg*mRHel-HA and r*Tg*mRHel-HA parasites cultured in the absence or presence of ATc. After reverse transcription, cDNA was amplified using a *coxI*-specific internal primer together with an adaptor-specific primer.

Analysis of the PCR products revealed three prominent bands (Fig 5A). In control samples, a single dominant band - designated band X - was detected. Upon depletion of *Tg*mRHel, the intensity of band X decreased markedly, while two larger amplification products - band Y and band Z - became increasingly abundant (Fig 5A).

**Fig 5.**
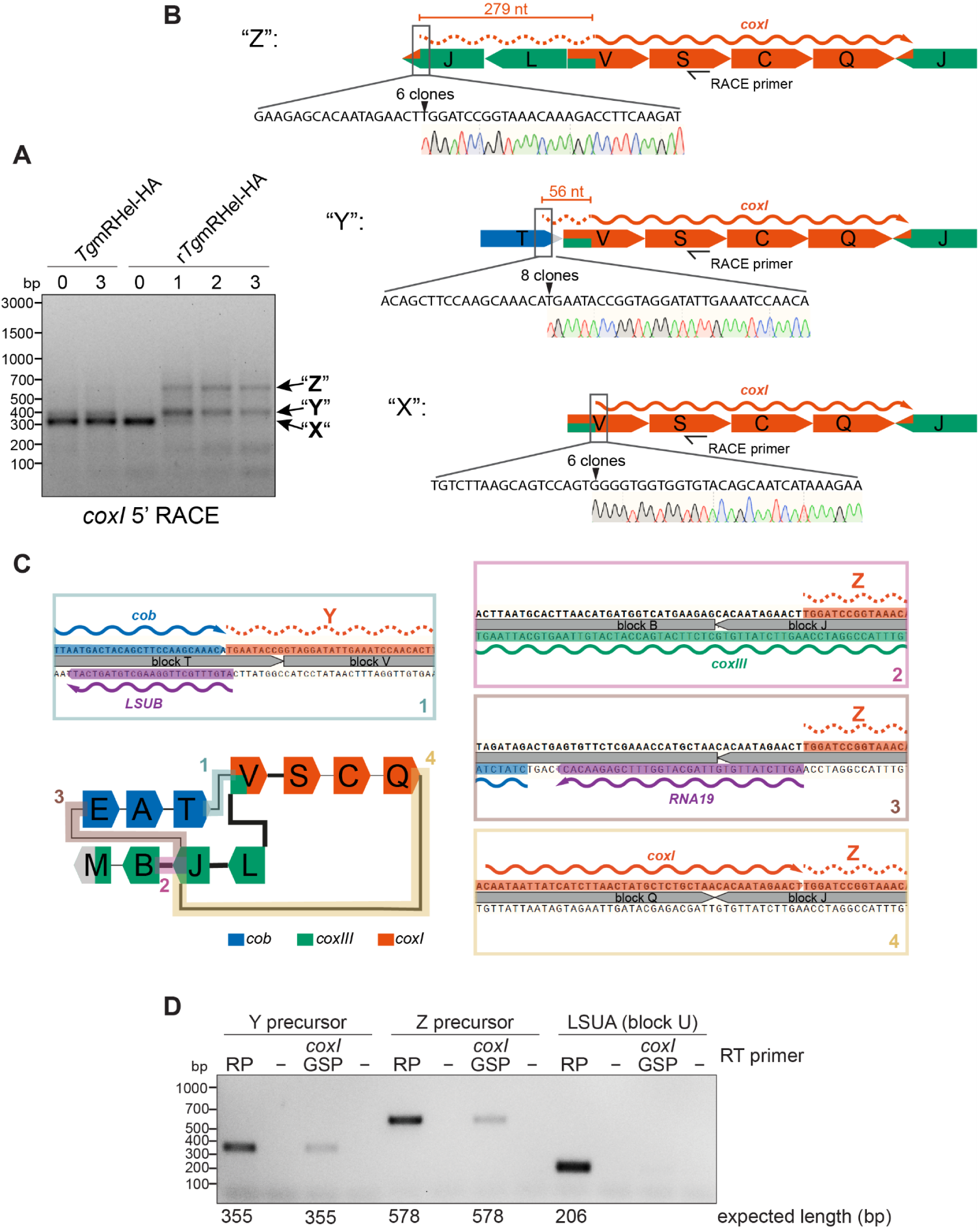
*coxI* precursor 5′-ends originate from different genomic contexts. A. Agarose gel analysis of 5′ RACE PCR products was performed to identify the 5′ ends of *coxI* transcripts. RACE was conducted on RNA extracted from r*Tg*mRHel-HA parasites cultured either in the absence or presence of ATc for 1–3 days, alongside the parental *Tg*mRHel-HA line grown with or without ATc for 3 days. Sequencing of the three major PCR products revealed that the two larger amplicons (designated as Y and Z) observed in *Tg*mRHel-depleted parasites represent precursor RNAs originating from different genomic blocks, while the smallest product (designated as X) corresponds to the mature *coxI* mRNA (see B). B. Schematic representation of *T. gondii* mitochondrial genomic sequence blocks encoding CoxI and alternative upstream genomic regions. Blocks are depicted as arrows indicating directionality, as published in [14], and are not drawn to scale. The gene-specific primer used for 5′ RACE is shown as a black one-sided arrow. Sequencing of *coxI* 5′ RACE PCR products, designated as X, Y, and Z in panel A, revealed distinct *coxI* transcripts illustrated by red wavy lines, with putative precursor regions shown as dotted regions. Zoom-ins indicate the precise locations of the transcript ends and the number of sequenced clones representing each end. Band X, representing the dominant *coxI* 5′ end in the parental line, maps to block V and likely corresponds to the mature 5′ end. Bands Y and Z accumulate following *Tg*mRHel depletion, with the end of Y mapping to block T, 56 nt upstream of the mature *coxI* end, and the end of Z mapping to block J, 279 nt upstream of the mature end in a distinct genomic context. C. Schematics visualizing the location of *coxI* precursor 5′ ends in different mitogenomic arrangements. Lower left panel: Visualization of genomic arrangements of mitochondrial sequence blocks encoding *coxI* (red), *cob* (blue), and *coxIII* (green), adapted from [13]. Additional combinations containing sequence blocks not relevant for *coxI* precursors were omitted for clarity. Colored block connections refer to the numbered, color-coded zoom-in windows. RNAs are shown as wavy arrows, with precursor regions indicated as dotted lines. The 3′ ends of *cob* and *coxI* are based on 3′ RACE experiments shown in S6 Fig; rRNAs are annotated according to [9,10,13]. (1) The 5′ end of precursor Y is immediately adjacent to the 3′ end of the *cob* mRNA and overlaps the 5′ end of the LSUB rRNA on the opposite strand by 1 nucleotide. (2–4) The 5′ end of precursor Z is located in block J, which occurs in three distinct genomic arrangements: (2) In the block J–B context, the precursor Z 5′ end is antisense to the *coxIII* mRNA.(3) In the block J–E context, the 5′ end lies adjacent to the 5′ end of RNA19 on the opposite strand. (4) In the block J–Q context, the 5′ end of *coxI* precursor Z lies directly adjacent to the *coxI* 3′ end. D Agarose gel electrophoresis of PCR products from reverse-transcription (RT)-PCR performed on TATi/Δku80 parasites, amplifying *coxI* precursor regions identified following *Tg*mRHel depletion. Reverse transcription was performed using random primers (“RP”) or gene-specific primers (“*coxI* GSP”) annealing to the *coxI* 3’ end. For both precursor Y and precursor Z, fragments of the expected size were amplified from cDNA generated with RPs or *coxI* GSPs and verified by Sanger Sequencing. The rRNA fragment LSUA encoded on block U was amplified as a control, showing amplification only from cDNA generated with RPs, supporting that the amplified precursor regions Y and Z originate from *coxI* transcripts. “-RT” = control without reverse transcriptase

To determine the origin of these transcripts, all three bands were cloned and sequenced. Sequencing of band X showed that its 5′ end maps within mitochondrial sequence block V, downstream of the previously reported 5′ end of the mature *coxI* mRNA [Fig 5B, 14]. The disappearance of band X upon *Tg*mRHel depletion mirrors the loss of the ∼1.5 kb mature *coxI* transcript observed in RNA gel blot analyses (Fig 4C). These results indicate that band X represents the mature *coxI* mRNA 5′ end.

Sequencing of the two larger products revealed that they correspond to longer *coxI* precursor transcripts with distinct upstream genomic contexts. The 5’ end of band Y maps to mitochondrial sequence block T, 56 nt upstream of the mature *coxI* 5′ end (Fig 5B). This region lies directly adjacent to the 3′ end of the *cob* transcript and the 5′ end of the LSUB rRNA fragment, which is encoded antisense to *cob* (Fig 5C, inset 1; S6 Fig). The precise coincidence of these transcript boundaries suggests that precursor Y is generated by cleavage at the *cob* 3′ end.

The 5’ end of band Z maps further upstream, within mitochondrial sequence block J, 279 nt upstream of the mature *coxI* 5′ end (Fig 5B). This sequence block occurs in multiple genomic configurations within the recombining mitochondrial genome of *T. gondii*. In one arrangement, block J is joined to block B, placing the precursor region antisense to the *coxIII* transcript (Fig 5C, inset 2). In another configuration, block J joins block E, positioning the precursor adjacent to the RNA19 rRNA fragment on the opposite strand (Fig 5C, inset 3). A third configuration links block J to block Q, where the 5′ end of the precursor lies directly adjacent to the 3′ end of another *coxI* copy (Fig 5C, inset 4). At present, it is not possible to determine which of these genomic contexts - or whether all of them - contribute to the Z precursor transcripts.

The observation that bands Y and Z accumulate specifically upon *Tg*mRHel depletion suggests that these transcripts represent processing intermediates that are normally converted into the mature *coxI* mRNA in a *Tg*mRHel-dependent process. Consistent with this interpretation, band Y was less abundant in untreated r*Tg*mRHel-HA parasites than in the parental line (Fig 5A). Because the r*Tg*mRHel-HA strain overexpresses *Tg*mRHel in the absence of ATc (Fig 1C), this reduction likely reflects increased processing efficiency under these conditions.

A remaining question was whether these precursor RNAs arise only upon loss of *Tg*mRHel or whether they are normally present at low levels in wild-type parasites. To address this, we performed RT-PCR using RNA from the parental TATi/Δku80 strain. Using primer pairs specific for the Y and Z precursor regions, amplification products of the expected sizes were detected when reverse transcription was performed with either random primers or *coxI*-specific primers and verified by Sanger sequencing (Fig 5D). These results demonstrate that the Y and Z precursor transcripts are also present in wild-type parasites, although at much lower abundance.

Together, these results indicate that the *coxI* mRNA can originate from multiple precursor transcripts generated from different genomic contexts within the recombining mitochondrial genome. *Tg*mRHel is required for the processing step that converts these heterogeneous precursor RNAs into a mature *coxI* transcript with a defined 5′ end.

## Discussion

### Loss of *Tg*mRHel impairs mitochondrial rRNA and *coxI* mRNA expression, Complex IV production, disrupting OxPhos and parasite growth

Mitochondrial gene expression in apicomplexan parasites such as *T. gondii* represents a highly specialized and poorly understood process. In this study, we identify and characterize *Tg*mRHel, a mitochondrially localized DEAD-box RNA helicase that is essential for parasite proliferation. Our findings show that *Tg*mRHel plays a central role in mitochondrial RNA metabolism, having multiple roles. It mediates the 5’-end processing of *coxI* mRNA and facilitates the accumulation of *coxIII* mRNA and mitochondrial rRNAs. These functions are critical for the maintenance of mitochondrial translation and thus for biogenesis of Complex III and IV of the electron transport chain (ETC). The defects in mitochondrial gene expression observed upon mRHel1 depletion culminate in a dramatic loss of oxygen consumption and the arrest of parasite proliferation, demonstrating the essential role of *Tg*mRHel in maintaining mitochondrial function.

It is currently unclear to what extent the various functions of *Tg*mRHel contribute individually to the strong reduction of Complex III and IV upon *Tg*mRHel depletion. The rapid decline in mature *coxI* and *coxIII* mRNAs likely impairs the synthesis of their encoded subunits, directly compromising Complex IV biogenesis. In parallel, *Tg*mRHel depletion leads to a marked reduction in mitochondrial rRNA fragments and a shift in their sedimentation profiles away from assembled ribosomal complexes. Together, these defects point to impaired mitochondrial translation, which likely exacerbates the reduction of Complex III and IV. Several RNA helicases, including DEAD-box family members such as Mrh4 and Mss116, are known to be essential for mitochondrial ribosome biogenesis in yeast [31,32] and humans [33,34] and for cytosolic ribosome assembly in other eukaryotes [35]. Most of these helicases also play additional roles in RNA metabolism, including RNA processing, surveillance, and decay [32,35–39] The multifunctional nature of DEAD-box helicases supports the view that *Tg*mRHel similarly operates at multiple levels of mitochondrial gene expression in *T. gondii*, coordinating both mRNA processing and ribosome biogenesis.

*Tg*mRHel shares structural features with the conserved DDX5/DDX17 subfamily of DEAD-box helicases. DDX5/DDX17 helicases play versatile roles throughout RNA metabolism [40]. While their ability to unwind RNA duplexes initially defined their enzymatic activity, these proteins also facilitate RNA strand annealing and remove RNA-bound proteins. Rather than acting in isolation, DEAD-box proteins typically function within large multi-protein complexes. Their functional diversity is believed to stem from interactions mediated by their structurally diverse N- and C-terminal extensions with various partner proteins. In this regard, it is noteworthy that *Tg*mRHel has an unusually long N-terminal intrinsically disordered region (IDR), which may provide structural flexibility or facilitate interactions with RNA or protein partners. Phylogenetic analysis indicates that *Tg*mRHel clusters within a group of apicomplexan- and dinoflagellate-specific DEAD-box helicases, suggesting functional specialization within this lineage. Dinoflagellate mitochondria share the highly reduced coding capacity of their genomes, the highly fragmented rRNAs of the ribosome, and complex coding gene architectures with apicomplexans. These features are likely linked with requirements for enhanced RNA processing abilities [13,14,41].

### RACE suggests an alternative *coxI* start codon

Previous phylogenetic analyses of the CoxI protein and reading frame could not unequivocally assign a start codon to the *coxI* mRNA [42], although a canonical AUG start codon has been suggested later [14, Fig S7]. In an alignment of related CoxI protein sequences, the sequence immediately downstream of the annotated start codon is not conserved, and conservation starts only at position 22 (S7 Fig). This is downstream of a possible alternative non-canonical start codon, which would correspond to amino acid position 17 in the alignment. If true, the usage of a non-canonical *coxI* start codon would be a feature consistent with observations in mitochondrial genomes of *P. falciparum* and dinoflagellates (Rehkopf et al. 2000, Feagin et al. 2012, Slamovits et al. 2007, Jackson et al. 2007, Waller and Jackson 2009), which form a sister group to apicomplexans.

### *Tg*mRHel-dependent processing of *coxI* mRNA from cryptic precursor RNAs

We observed the accumulation of aberrant *coxI* precursor transcripts in *Tg*mRHel-depleted parasites and the concurrent loss of the mature *coxI* mRNA. This suggests that *Tg*mRHel acts post-transcriptionally to process *coxI* precursor RNAs into mature transcripts. RACE analyses demonstrate that these precursor transcripts arise from distinct genomic contexts, reflecting the modular, recombining nature of the *T. gondii* mitochondrial genome [13,14]. In addition, *Tg*mRHel is also required for the accumulation of *coxIII*, but there is no evidence for the accumulation of precursor RNAs. The third mitochondrial mRNA, *cob*, is largely unaffected in the mutant.

Although our data do not directly reveal the mechanism by which *Tg*mRHel stabilizes transcripts or processes precursor RNAs, they do allow us to infer steps involved in the processing of the *coxI* 5′-end. Notably, the precise positioning of the *coxI* precursor 5′-end adjacent to the 3′- and 5′-ends of *cob* and *coxI* mRNAs, and the RNA19 rRNA fragment suggests that these ends are generated by highly specific nucleolytic events (Fig 5). It further suggests the existence of long precursor RNAs that encompass additional sequences, for example *coxI* along with either *cob* or a second *coxI* copy, depending on the genomic context. The processing of such long precursors likely involves endonucleolytic cleavage events. The finding that overexpression of *Tg*mRHel leads to loss to the “Y” precursor seen in the parental line further supports the idea that the helicase fosters processing of precursors to mature *coxI*.

How might this processing occur? One possibility is that the region to be trimmed exists initially as a double-stranded RNA (dsRNA), which must be unwound to allow nuclease access. This could involve an endonuclease, although it’s unclear how such an enzyme would be targeted specifically to the mature *coxI* 5′-end. RAP proteins have been suggested to have endonuclease activity and might be candidates here [43–46]. An alternative model is that *Tg*mRHel facilitates the activity of a 5′→3′ exonuclease that degrades the *coxI* precursor RNA from the 5′-end. In yeast and metazoan mitochondria, 3′→5′ exonucleolytic degradation is mediated by the mitochondrial degradosome, which includes the exonuclease PNPase and the RNA helicase Suv3p [47–49]. Within this complex, Suv3p unwinds double-stranded RNA structures to enable PNPase-mediated exonucleolytic activity from the 3′-end, supporting transcript maturation and quality control and preventing the accumulation of dsRNA arising from non-specific mitochondrial antisense transcription [50,51]. 5′→3′ processing is mediated in yeast by Pet127 and a link to the Suv3p helicase has been suggested based on rescue of a Suv3p mutant by overexpressing Pet127 [52,53]. Whether *Tg*mRHel fulfils similar functions as Suv3p will require a biochemical characterization of its catalytic activity and identification of protein and RNA interaction partners. Also, it is at present unclear why the exonucleolytic leader removal stops at the specific 5’-end observed for *coxI* here. RNA binding proteins like FASTKD proteins have been shown to act in the formation of mammalian mitochondrial transcript ends that are not generated by tRNA processing [54,55]. FASTKD proteins belong to a larger family of helical repeat RNA binding proteins that can act as highly specific roadblocks against organellar exonucleases [55–57], or, if they contain a RAP domain, can also function as endonucleases [45]. In Apicomplexa, this family has expanded and is known as heptatricopeptide repeat (HPR) proteins [57]. Two HPR proteins with RAP domains have been shown to directly bind to rRNAs in *Plasmodium falciparum* [44]. An intriguing question is whether such HPR or HPR–RAP proteins cooperate with *Tg*mRHel in generating the *coxI* 5’-end in *T. gondii*.

### mRHel unifies mRNA production from different genome-based precursor RNAs in T. gondii

Transcript diversity arises mostly through RNA-level processes, including alternative splicing, RNA editing, endonucleolytic cleavage, and exonucleolytic trimming. DNA contributes to transcript variation primarily through differential promoter usage and alternative polyadenylation sites, which can influence the length and structure of the primary transcript. However, in the highly fragmented and recombined mitogenome of *T. gondii*, transcript complexity originates already at the DNA level. The three mRNAs under investigation are encoded in multiple genomic contexts. As a result, transcripts can initiate and terminate at different genomic sequence blocks, giving rise to a variety of precursor RNAs with distinct 5′ and 3′ untranslated regions. Our 5′ RACE analyses demonstrate that transcripts arise from alternative combinations of genomic blocks. Yet, Northern blot analysis reveals that these precursor RNAs are nearly undetectable in control lines. Only a single precursor RNA species is identified for *coxI*, with no detectable precursors for *cob* or *coxIII*, suggesting that precursor RNAs are rapidly and efficiently processed into mature mRNAs. Thus, regardless of their genomic origin, precursor RNAs appear to undergo non-specific removal of their variable leader sequences. Speculatively, the peculiar organization of the *T. gondii* mitochondrial genome has led to the need for an adaptable RNA processing system capable of handling precursors with diverse structures and origins, although reverse explanations - that enhanced RNA processing allowed more diverse genomic configurations - cannot be excluded. Our analysis demonstrates that *Tg*mRHel is a component of this RNA processing machinery. Depletion of *Tg*mRHel results in a marked reduction of *coxI* mRNA levels within 24 hours, accompanied by a decrease in Complex IV abundance and ensuing strong growth defect, highlighting the essential role of leader removal. In conclusion, our findings reveal a novel situation in which genome rearrangements generate transcript diversity, necessitating a unifying RNA processing activity to produce mature mRNAs. The involvement of *Tg*mRHel in non-specific leader removal provides an elegant solution to this challenge. From an evolutionary perspective, this processing flexibility may buffer the effects of frequent genomic recombination in *T. gondii* by rescuing potentially deleterious precursor configurations, while also preserving - or even expanding - the functional diversity generated by advantageous recombination events.

Recombination-driven generation of gene isoforms also occurs in many ciliates. For example, *Oxytricha trifallax* possesses a highly scrambled genome in its micronucleus - the germline nucleus used during sexual reproduction - while its somatic macronucleus is formed by extensive DNA rearrangement, yielding thousands of gene-sized chromosomes that are actively transcribed. Variability in recombination during macronuclear development can produce alternative genomic configurations that encode the same ORF but differ in their UTRs and in the coding sequence itself (Swart et al. 2013; Klobutcher et al. 1988; Chen et al. 2015). Whether such recombination-based isoforms pose similar challenges for gene expression as in the *T. gondii* mitogenome - and whether comparable solutions have evolved - remains an intriguing question.

## Materials and Methods

### Parasite culture and Plaque assays

*T. gondii* was cultured in mycoplasma-free human foreskin fibroblasts (HFFs; SCRC-1041™ ATCC). Parasites were grown in DMEM with 1% fetal bovine serum and antibiotics. When required, ATc was added at 0.5 µg/ml. Plaque assays followed the method of van Dooren et al. [58].

### Generation of modified parasites

All transgenic parasite lines were generated using the CRISPR/Cas9 system as previously described [59]. Single guide RNAs (sgRNAs) were selected using CHOPCHOP [60 https://chopchop.cbu.uib.no/], based on predicted on-target efficiency and minimal off-target effects. Primers used in all the steps described are listed in S2 Table.

The parental parasite line, *Tg*mRHel-HA, was generated by C-terminally tagging *Tg*mRHel. We modified pSAG1::Cas9-U6::sgUPRT [Addgene plasmid # 54467; ,59] by Q5 site-directed mutagenesis (NEB) using a gene specific primer *Tg*mRHel 3’ tag CRISPR fwd and the universal CRISPR reverse primer listed in S2 Table. This generated a sgRNA-expressing vector targeting the genomic region near the stop codon of *Tg*mRHel. The vector also encoded GFP-tagged Cas9. It was co-transfected into TATi/Δku80 parasite strain [61] along with a PCR-generated donor template containing the HA-tag followed by a DHFR selectable marker. A donor template was amplified from a plasmid plinker-3xHA-DHFR (Tetzlaff et al., 2024) using primers *Tg*mRHel 3’ tag fwd and *Tg*mRHel 3’ tag rev with 50 bp of homology to the regions upstream and downstream of the stop codon of *Tg*mRHel (S2 Table). Following transfection, parasites were selected on pyrimethamine, and clones were obtained by limiting dilution. Successful integration of the HA tag was confirmed by PCR [62] with genotyping primers listed in S2 Table.

To generate anhydrotetracycline (ATc)-regulated version of the *Tg*mRHel-HA line (named r*Tg*mRHel-HA) we replaced the endogenous promoter of *Tg*mRHel-HA with an ATc-regulated promoter as described previously [61,63]. A similar CRISPR/Cas9 approach as above was used with primers listed in S2 Table. We prepared a sgRNA-expressing vector targeting the genomic region around the start codon of *Tg*mRHel and co-transfected it together with a donor template. A donor template containing chloramphenicol resistance marker followed by an ATc-regulated t7s4 promoter was PCR-amplified from a plasmid pPR2-HA3-Floxed-CAT [a modified version of pPR2-HA3; ,63] using primers (S2 Table) with 50 bp of homology to the regions upstream and downstream of the start codon of *Tg*mRHel. After transfection, parasites were selected on chloramphenicol, clones were obtained by limiting dilution and screened for successful promoter replacement by PCR with genotyping primers listed in S2 Table.

For complementation of the r*Tg*mRHel-HA line with constitutively expressed mRHel-Ty1, the mRHel ORF was synthesized (Twist Bioscience) and integrated into pUDT-Ty1 [30]. The region containing the alpha-tubulin promoter followed by the mRHel ORF fused to a 3×Ty1 tag sequence was amplified from the vector using primers with overhangs homologous to the *T. gondii* UPRT locus (S2 Table). The PCR product was co-transfected together with the pSAG1::Cas9-U6::sgUPRT vector expressing an sgRNA targeting the UPRT locus into the r*Tg*mRHel-HA line. Selection was performed with 10 μM 5-fluorodeoxyuridine (FUDR), parasites were cloned by limiting dilution, and clones PCR-screened for successful mRHel-Ty1 integration.

All subsequent lines were derived from r*Tg*mRHel-HA. They were generated as described previously [29] by introducing a FLAG-tag fusion to the following target genes: *Tg*SdhB, *Tg*MppA, *Tg*ApiCox25 and *Tg*ATPb. Existing sgRNA-expressing vectors targeting the genomic region near the stop codon of *Tg*SdhB, *Tg*MppA and *Tg*ApiCox25 (Seidi et al., 2018; Hayward et al., 2021; Leonard et al., 2023) were used. A donor template with FLAG-tag was amplified by PCR from a template synthesized as a gBlock [IDT; 29] using gene-specific primers containing 50 bp of homology to the regions flanking the stop codon of the *goi* (S2 Table). For each of the target genes, a separate transfection into the r*Tg*mRHel-HA parasite line was carried out. Three days post-transfection GFP-positive parasites were sorted and cloned by flow cytometry. Successful integration was confirmed by PCR as described [62] using genotyping primers listed in S2 Table.

### Analysis of respiratory activity and glycolysis

A Seahorse XFe96 extracellular flux analysis was performed as described to determine respiratory and glycolytic activity [28].

### SDS-PAGE, BN-PAGE and western blotting

SDS-PAGE and western blotting were performed as described [13,58] using the following antibodies: mouse anti-FLAG clone M2 (1:500, Sigma, F3165), rat anti-HA High Affinity clone 3F10 (1:200-1:500, Roche, ROAHAHA), and rabbit anti-*Tg*Tom40 [1:2000, 64]. Horseradish peroxidase (HRP)-conjugated secondary antibodies goat anti-rat IgG (1:10000, Abcam, catalog number ab97057), goat anti-mouse IgG (1:5000, Abcam, catalog number ab6789) and goat anti-rabbit IgG (1:10000, Abcam, catalog number ab97051) were used.

Blue native PAGE followed an established protocol [64]. Blots were probed with the following primary antibodies: mouse anti-FLAG clone M2 (1:800, Sigma, F3165). HRP-conjugated goat anti-mouse IgG (Abcam, catalog number ab6789) secondary antibodies were used at 1:5000.

### Polysome analysis and sRNA gel blot

Sucrose gradient analysis and RNA gel blot analysis of small mitochondrial RNA of mRNAs were performed as described [13] with some modifications. Specifically, the benzonase step was omitted from the organelle enrichment procedure. Instead, the pellet after 0.05% digitonin treatment and centrifugation was frozen at -80°C for at least overnight, and after thawing, subjected to lysis under conditions optimized for mitoribosome stability (30 mM MgCl2, 100 mM KCl).

### RACE

For 5’ RACE total RNA from *T. gondii* was ligated to a small RNA oligo (Rumsh) using T4 RNA Ligase 1 (NEB) following the manufacturer’s protocol. Reverse transcription was performed with ProtoScript® II Reverse Transcriptase (NEB) and random primers. cDNA was amplified by PCR using gene-specific primers and an oligo-specific primer (Rumsh2) with Taq DNA Polymerase (Roboklon). The PCR product was excised from the gel, purified, and cloned using the CloneJET PCR Cloning Kit (NEB) according to the manufacturer’s instructions. Plasmids from transformed *E*. *coli* were isolated and sequenced by Sanger sequencing (Azenta Life Sciences).

For 3’ RACE, total RNA was reverse transcribed using an oligo(dT) primer containing a 3’ linker sequence (3’ RACE RT primer). The PCR was performed using gene-specific primers in combination with a linker-specific primer (3’ RACE linker-specific primer). PCR products were gel-extracted, cloned, and analyzed as described for 5’ RACE.

### mRNA gel blot hybridization analysis

mRNA gel blot hybridizations were performed as described previously [65].

### Immunofluorescence/Fluorescence microscopy assays

Immunofluorescence assays (IFAs) were performed as described by van Dooren et al. [58]. Primary antibodies included rat anti-HA High Affinity clone 3F10 (1:500, Roche, ROAHAHA), mouse anti-FLAG M2 (1:250-1:500, Sigma, F3165), and rabbit anti-*Tg*Tom40 [1:2000, 64] . Secondary antibodies used were goat anti-rat Alexa Fluor 488 (1:500, Thermo Fisher Scientific, A-11006), goat anti-mouse Alexa Fluor 488 (1:500, Thermo Fisher Scientific, A-11029), and goat anti-rabbit Alexa Fluor 594 (1:500, Thermo Fisher Scientific, A11012). Images were acquired on a DeltaVision Elite deconvolution microscope (GE Healthcare) with a 100X UPlanSApo objective lens (NA 1.40). They were deconvolved using SoftWoRx Suite 2.0 software and contrast and brightness were adjusted in FIJI/ImageJ. Images were then processed with Adobe Illustrator.

## Acknowledgements

We thank Michael Devoy and Harpreet Vohra (John Curtin School of Medical Research, ANU) for performing cell sorting and Edwin Tjhin (van Dooren Lab, ANU) for the preparation of sgRNA-expressing vector targeting the *Tg*ATPb.

---

**Fig S1.**
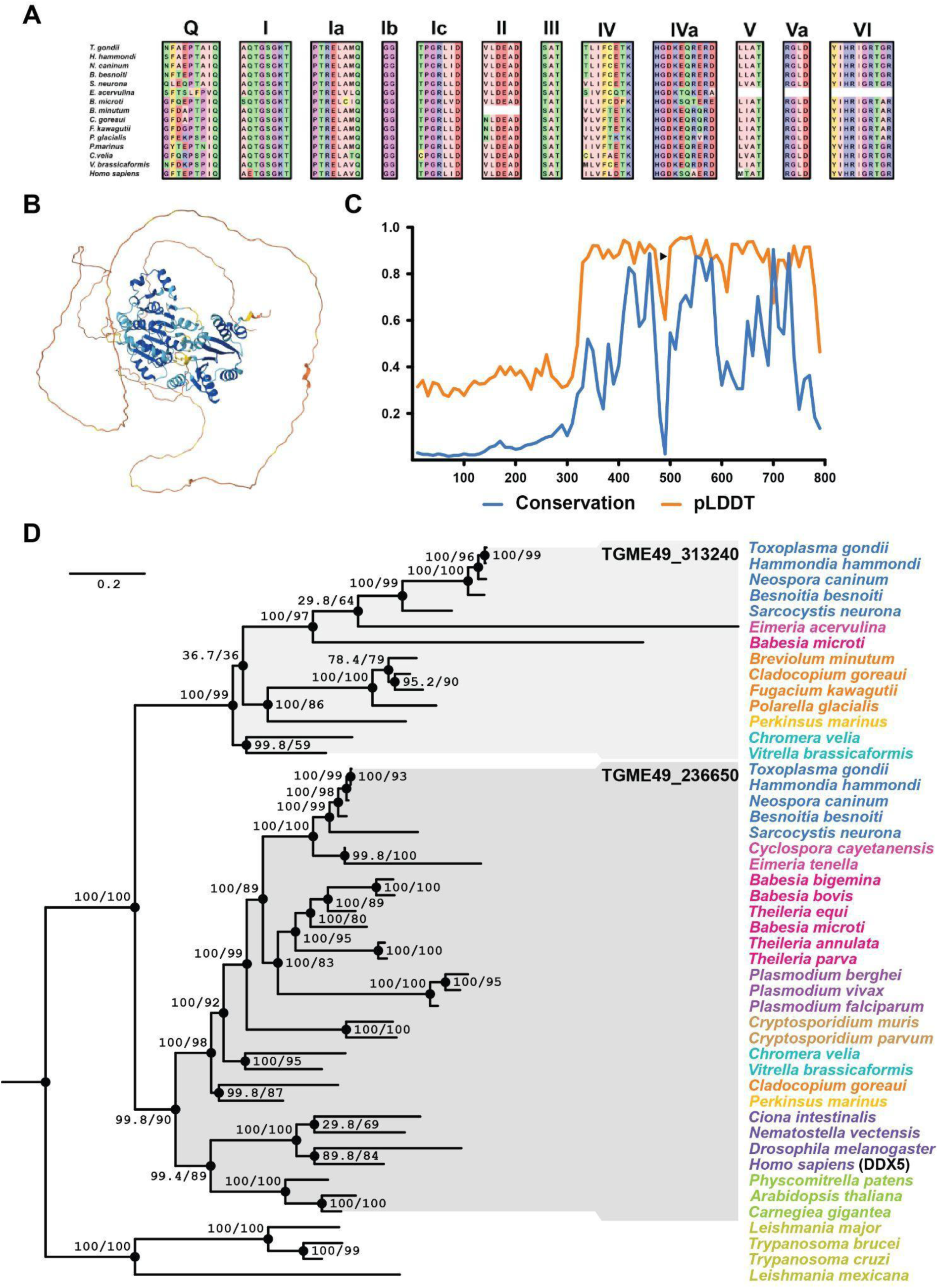
Structural and phylogenetic analysis of *Tg*mRHel (TGGT1_313240) A. Protein sequences from selected eukaryotic species were aligned to illustrate the conserved motifs characteristic of DEAD-box helicases. The alignment was initiated using the *Toxoplasma gondii* sequence, with homologs from other organisms identified by BLAST searches. Conserved sequence motifs (Q, I, Ia, Ib, Ic, II, III, IV, IVa, V, Va, VI) are boxed according to their known positions in DEAD-box helicase structures. Motif II (Walker B) contains the name-giving amino acids DEAD. Sequences outside the conserved motifs (N- and C-terminal extensions) are not shown. B. Structural prediction was generated using AlphaFold for the full-length *Tg*mRHel protein. The model highlights the conserved helicase core domain (shown in blue), which exhibits the characteristic fold of the DEAD/H-box family, including canonical RecA-like domains. The extensive N-terminal region is predicted to be intrinsically disordered, as indicated by its lack of defined secondary structure and low confidence scores (colored from yellow to red). This disordered region is hypothesized to mediate regulatory or localization functions. This supplemental figure complements the schematic domain organization shown in the main text (Fig 1A), providing structural context for both the conserved helicase motor and the unique extended N-terminal region of *Tg*mRHel. C. Predicted disorder profile of *Tg*mRHel generated using the flDPnn server (Hu et al. 2021). Disorder probability is plotted against amino acid position; higher scores reflect increased likelihood of intrinsic disorder. Disorder and conservation profiles of *Tg*mRHel obtained from the alphafold3 predicted structure and the alignment of the helicases shown in panel D, respectively. pLDDT was used as a metric to assess the disorder of *Tg*mRHel with lower values indicating propensity to disorder. Conservation was calculated as the proportion of residues that were identical to those of *Tg*mRHel at each position. D. Phylogenetic analysis of *Tg*mRHel and related DEAD/H-box RNA helicases. Sequences from reciprocal best hits of *Tg*mRHel in 37 organisms were aligned using mafft, from the aligned sequences, a ML phylogenetic tree was constructed with iqtree2 under the LG+I+R4 substitution model. The tree indicates that *Tg*mRHel originated from an ancient gene duplication event with DDX5 orthologues including TGME49_236650. Human protein DDX5 is shown.

**Fig S2.**
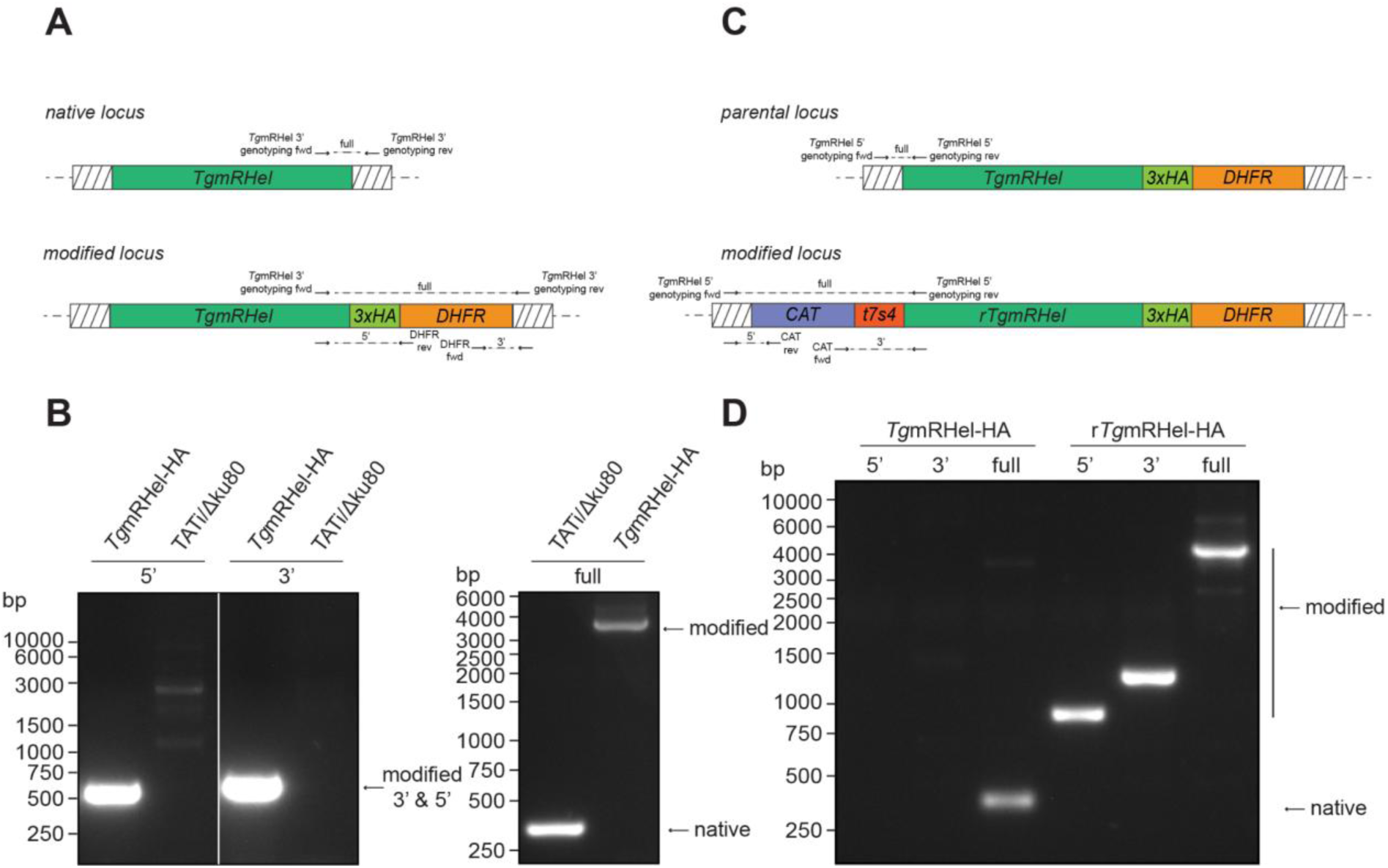
Tagging and knock-down approach for *Tg*mRHel (TGGT1_313240) A. Top panels: Schematic showing the 3′ replacement strategy used to insert a triple HA tag into the *Tg*mRHel locus. The top panel illustrates the native *Tg*mRHel locus, while the bottom panel shows the locus after insertion of the HA tag at the 3′ end of the open reading frame together with the DHFR selectable marker. Primer binding sites used to screen for successful integration are indicated. B. PCR analysis of genomic DNA from putative *Tg*mRHel-HA parasite strain using the primer pairs shown in (A). Parental TATi/Δku80 genomic DNA was included as a control. The results indicate successful HA tag integration. C. Top panel: Diagram showing the insertion of an anhydrotetracycline (ATc)-regulatable teto7/sag4 (t7s4) promoter, along with a chloramphenicol acetyltransferase (CAT) resistance cassette, upstream of the start codon of the *Tg*mRHel gene, which had previously been modified to express a C-terminal triple HA tag (see A). Primer binding sites used to screen for promoter integration for both native and modified loci are indicated. D. PCR analysis of genomic DNA from putative ATc-regulatable r*Tg*mRHel-HA clones using the primers shown in (C) to assess for integration. Parental (*Tg*mRHel-HA) parasite line genomic DNA was included as a control. Results indicate successful integration.

**Fig S3.**
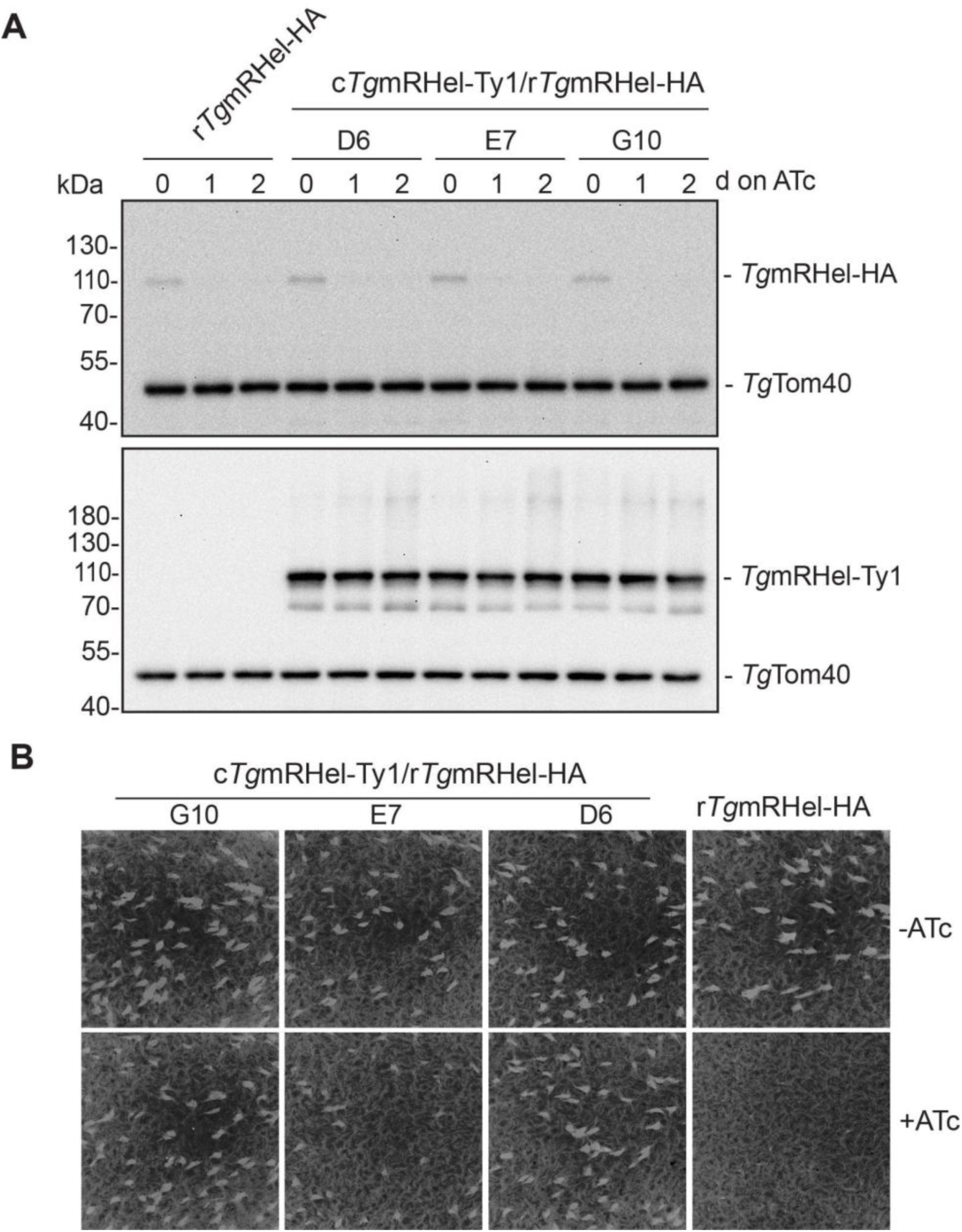
Complementation of r*Tg*mRHel-HA with constitutively-expressed *Tg*mRHel-Ty1 restores parasite proliferation. A. Western blots of proteins extracted from complementation parasite lines c*Tg*mRHel-Ty1/r*Tg*mRHel-HA (three independent clones G10, E7 and D6) cultured next to the parental line r*Tg*mRHel-HA in the absence of ATc or in the presence of ATc for one or two days. Samples were separated by SDS-PAGE, and probed with anti-HA and anti-*Tg*Tom40 (loading control) antibodies (upper panel). The same samples were analyzed on a separate membrane with anti-Ty1, and anti-*Tg*Tom40 antibodies (lower panel). B. Plaque assays measuring proliferation of r*Tg*mRHel-HA and c*Tg*mRHel-Ty1/r*Tg*mRHel-HA parasites cultured in the absence or presence of ATc for 8 days.

**Fig S4.**
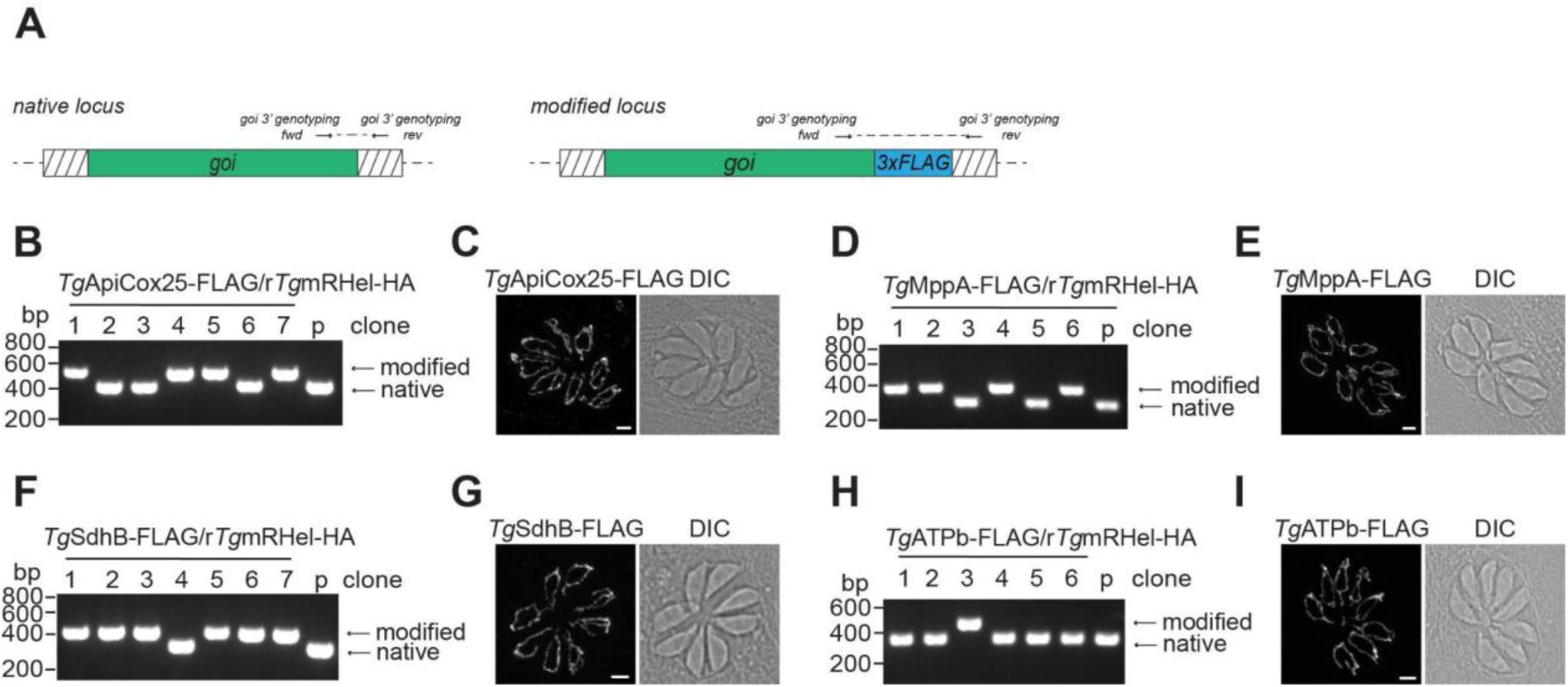
FLAG-tagged versions of nuclear-encoded OxPhos proteins *Tg*SdhB, *Tg*MppA, *Tg*ApiCox25 and *Tg*ATPb were integrated into r*Tg*mRHel-HA and are expressed. A. Top panels: Schematic showing the 3′ replacement strategy used to insert a triple FLAG tag into the loci of four nuclear-encoded subunits of OxPhos in an r*Tg*mRHel-HA background. Primer binding sites used to screen for successful integration are indicated and listed in S2 Table. Selection for integration was performed by FACS, based on GFP fluorescence from the co-transfected Cas9-GFP expression vector. B. PCR analysis of genomic DNA from putative *Tg*ApiCox25-FLAG/r*Tg*mRHel-HA clones using the primer pairs listed in S2 Table. The parental line r*Tg*mRHel-HA (p) genomic DNA was included as a control. The results indicate successful FLAG tag integration in clones 1, 4, 5 and 7. C. Immunofluorescence assay of eight-cells vacuole of *Tg*ApiCox25-FLAG/r*Tg*mRHel-HA parasites probed with anti-FLAG (white). Scale bar is 2 µm. D. PCR analysis analogous to panel (B) for *Tg*MppA-FLAG/r*Tg*mRHel-HA clones. The results indicate successful FLAG tag integration in clones 1, 2, 4 and 6. E. As in (C) for *Tg*MppA-FLAG/r*Tg*mRHel-HA. F. PCR analysis analogous to panel (B) for *Tg*SdhB-FLAG/r*Tg*mRHel-HA clones. The results indicate successful FLAG tag integration in clones 1, 2, 3, 5, 6 and 7. G. As in (C) for *Tg*SdhB-FLAG/r*Tg*mRHel-HA. H. PCR analysis analogous to panel (B) for *Tg*ATPb-FLAG/r*Tg*mRHel-HA clones. The results indicate successful FLAG tag integration in clone 3. I. As in (C) for *Tg*ATPb-FLAG/r*Tg*mRHel-HA.

**Fig S5.**
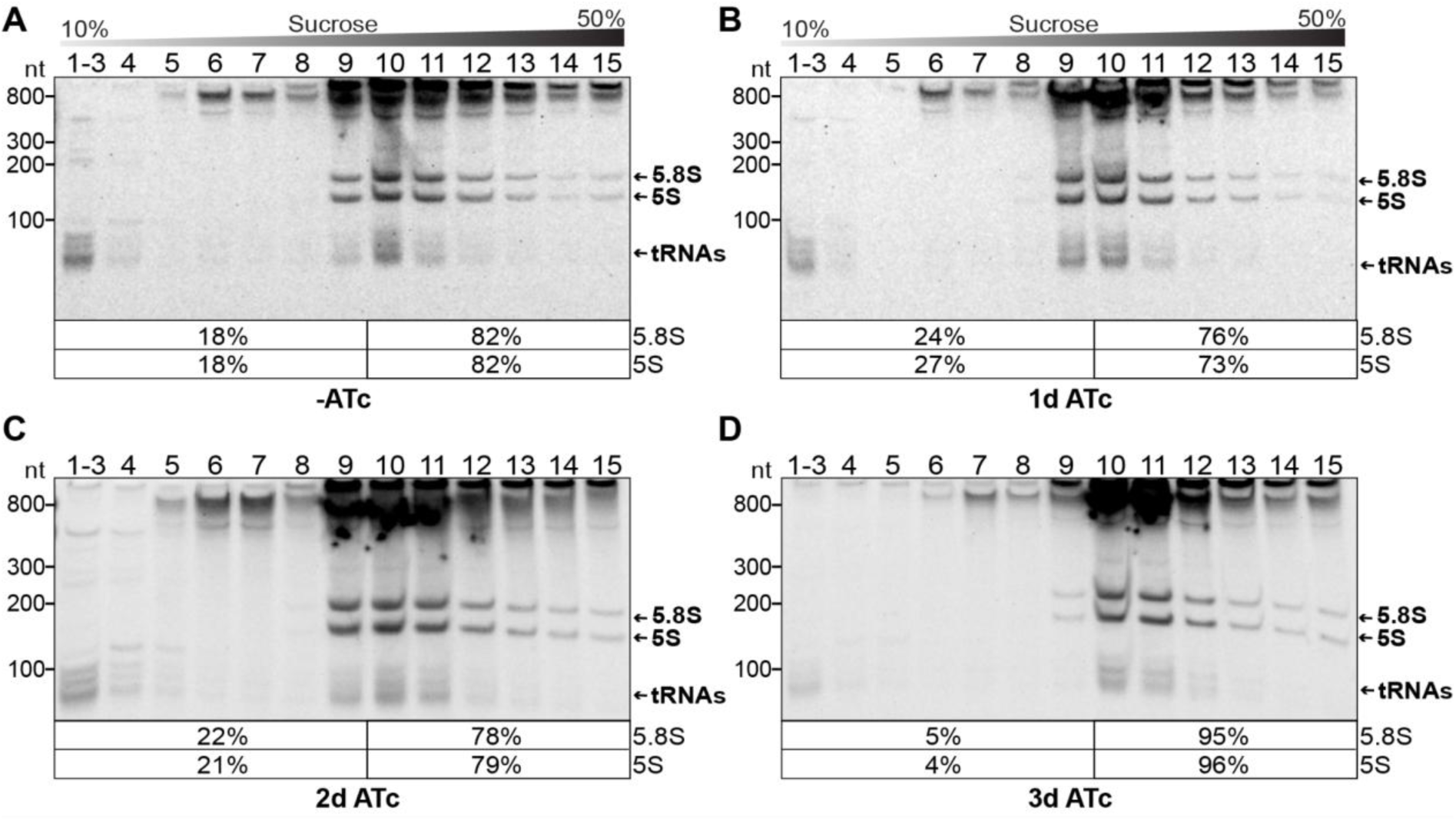
Ethidium bromide stain of PAGE gels after sucrose gradient fractionation of RNA-protein complexes in parasites depleted of *Tg*mRHel. Crude organelle-enriched extracts from *T. gondii* r*Tg*mRHel parasites either untreated (A), or treated with ATc for 1 (B), 2 (C) or 3 days (D) were separated on denaturing PAGE gels. These were stained with ethidium bromide to visualize abundant cytosolic rRNAs and tRNAs to control for equal loading. Atop the gel images, a schematic representation illustrates the sucrose density gradient. Percentages in square brackets below each blot show the proportion of 5.8S rRNA and 5S rRNA signal detected in fractions 1–9 relative to fractions 10–15.

**Fig S6.**
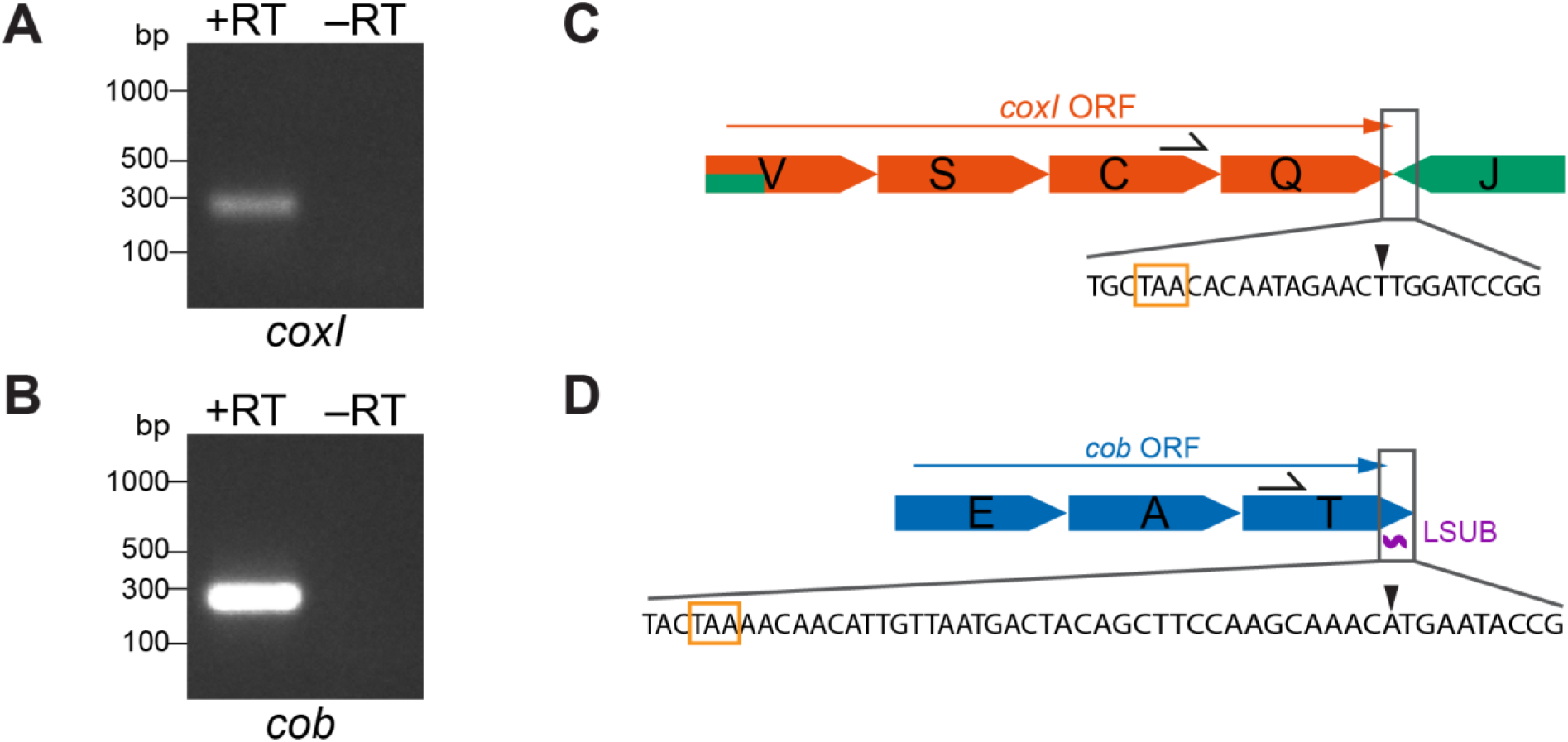
Identification of the *T. gondii coxI* and *cob* 3’ mRNA termini using 3’ RACE. A. PCR products of *coxI* 3’ RACE were analyzed by agarose gel electrophoresis. +RT and −RT denotes presence and absence of reverse transcriptase during cDNA synthesis, respectively. The amplicon was gel-extracted, cloned and analyzed by Sanger sequencing, with results shown in panel C. B. Same as panel A, but for *cob* 3’ RACE. C. Schematic representation of *T. gondii* mitogenomic sequence blocks encoding CoxI, with the proposed open reading frame (ORF) (Namasivayam et al. 2021) indicated by a red arrow. The location of the gene-specific RACE primer is shown by a black one-sided arrow. The zoom-in visualizes the 3’ region of the ORF downstream of the proposed stop codon (orange frame), with the 3’ end detected by RACE marked by a black triangle. The end was defined as the last nucleotide aligning to the reference sequence, followed by a poly(A) tail and the 3’ linker sequence. D. Same as in panel B, but for Cob. The LSUB rRNA fragment encoded on the opposite strand is shown as a purple wavy line.

**Fig S7.**
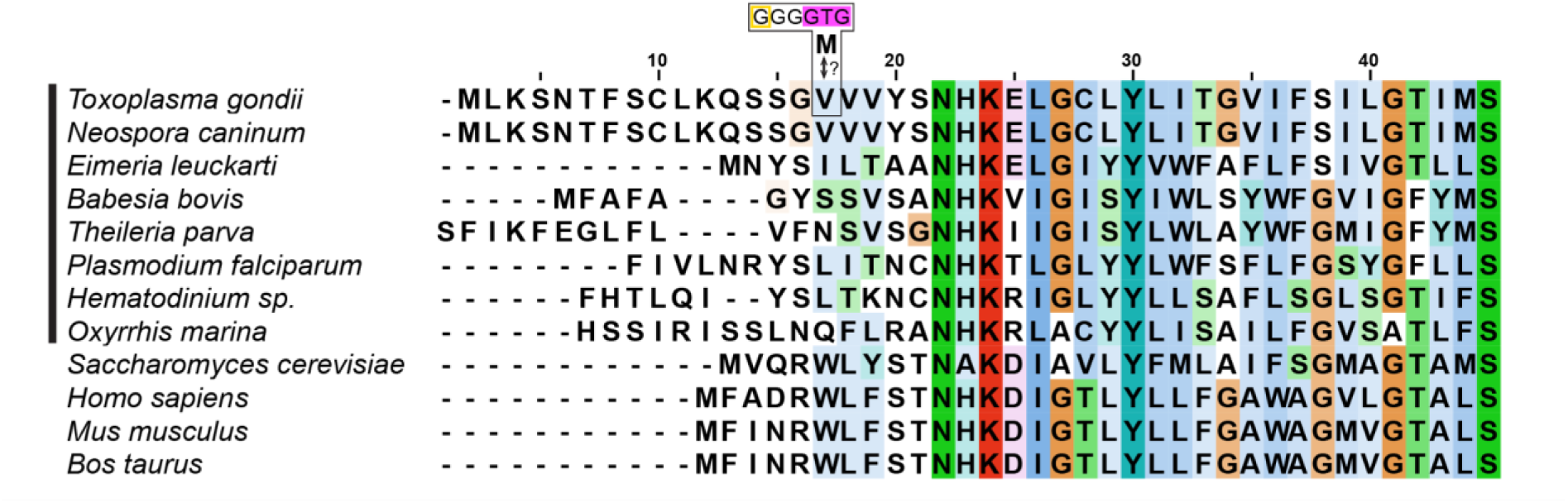
Multiple sequence alignment of CoxI N-terminal regions across different species. CoxI amino acid sequences of species representing the alveolate clade (marked by a black line next to the taxa names) and selected eukaryotic model organisms were aligned using Clustal Omega. Amino acids are colored according to the clustal coloring scheme with consensus-based intensity shading. Sequence conservation starts at the Asparagine (N) at position 22. In *T. gondii*, an in-frame GTG (shaded in pink), corresponding to amino acid position 17 in the alignment, may represent the non-canonical CoxI start codon. This potential start-codon is located upstream of the conserved region and downstream of the mRNA 5’ end identified by 5’ RACE (encircled in yellow; see Fig 5B), but downstream of the previously suggested canonical ATG shown here at the start of the *Toxoplasma* sequence (“<u>M</u>LK”; (Namasivayam et al. 2021)).

